# Hematological and genetic basis of clinical heterogeneity of HemoglobinE/β-thalassemia in Bangladesh

**DOI:** 10.64898/2026.02.22.707245

**Authors:** Farjana Akther Noor, Mohabbat Hossain, Suprovath Kumar Sarker, Khalid Arafath, Subarna Sayed Ety, Jeba Atkia Maisha, Mohammad Al Mahmud-Un-Nabi, Nusrat Sultana, Golam Sarower Bhuyan, A K M Ekramul Hossain, Waqar Ahmed Khan, Hossain Uddin Shekhar, Firdausi Qadri, Kaiissar Mannoor

## Abstract

Patients with HbE/β-thalassemia inheriting the same β-globin mutations display varied clinical manifestations, the mechanism of which is only partially known. The study aimed to decipher the heterogenous basis of HbE/β-thalassemia patients in more details by focusing on both hematological and genetic modifiers influencing the disease severity, which included-(i) HbF and HbE levels using Hb electrophoresis, (ii) β-thalassemia mutations, (iii) αααanti3.7triplication using Gap-PCR, (iv) individual and cumulative effects of HbF-inducing SNPs in 4 major modifier genes, namely HBG2, BCL11A, HBSB1L_MYB intergenic-region, and HBBP1 which were genotyped using DNA sequencing and Real-time PCR-HRM methods. Accordingly, 130 diagnosed Bangladeshi patients with HbE/β-thalassemia were enrolled and categorized as mild, moderate, and severe as per Mahidol scoring system. c.79G>A+IVS1_5G>C was the most predominant (73.8% of total) mutation pair across all the 3 severity groups, indicating secondary modifiers might influence the severity. Our study found both HbF and HbE protective to HbE/β-thalassemia, as both were inversely related to the severity score (HbF: p<0.0001/r=-0.55; HbE: p<0.0001/r=-0.56). Four SNPs-XmnI-Gγ, rs2071348 (HBBP1), rs489544 and rs28384513 (HBS1L_MYB) showed significant association with the elevated HbF levels (p=0.005, 0.0001, 0.0001, 0.004 respectively). The multivariate analysis showed that the risk genotypes with single or combination of 2, 3,and 4 SNPs showed gradually increased risk [Odd Ratio (95%CI)= 2.51, 5.47, 19.5, 39.0, respectively] of less severe phenotype, suggesting that these linked SNP variants had a cumulative effect on both HbF level and clinical severity score. However, low HbE level and copresence of αααanti3.7triplication were found to nullify the ameliorating effect of multiple SNPs.

## Introduction

Hemoglobin (Hb)E/β-thalassemia is one of the most serious inherited monogenic blood disorders worldwide^1, 2^. The patients with HbE/β-thalassemia have coinheritance of a β-trait from one parent and HbE-trait from the other. Clinical manifestations of HbE/β-thalassemia are highly heterogeneous which may be in the form of (1) mild with occasional blood transfusion or non-transfusion-dependent (NTD) form, (2) moderately severe-with blood transfusion frequency of more than the NTD group and less than the severe group, and (3) severe group-suffers from many severe complication and requires frequent blood transfusion^3^. However, ultimately mild patients also require regular care and adhere to the medications and iron chelation therapy^4^. Clinical heterogeneity of the disease, which is still only partially known, has made better management of these patients difficult^5^.

Though the causative mutations in the β-globin gene is the primary modifier of HbE/β-thalassemia, patients inheriting the same β-globin gene mutations are found to display heterogeneous clinical manifestations^6, 7^. To delineate the mechanism of clinical heterogeneity of the disease, the majority of the previous studies had focused on HbF-inducing *XmnI* polymorphism, single-nucleotide polymorphisms (SNPs) in the *BCL11A* gene, and other genetic loci like HBS1L-MYB, HBBP1, KLF1 etc^6, 8–12^. Production of high levels of HbF have been known to cause mild form of the disease^13^. Gamma-globin chain is capable of reducing the disease pathology by binding with excess unbound α-globin chain and preventing them from aggregation to cause apoptosis of nucleated erythrocytes through formation of HbF^14^. Although HbE plays dual roles to perform disease amelioration-(1) by neutralization of excess unbound α globin chains and (2) transporting O_2_ to the tissues to fulfil the O_2_ demand, only few reports had demonstrated a disease ameliorating role of HbE^15, 16^. Furthermore, co-inheritance of α-thalassemia produces a less severe disease minimizing the excess of α-globin chains while the α-globin gene triplication may produce severe phenotype when coinherited with HbE/β-thalassemia^17, 18^.

However, previous reports regarding HbF inducing SNPs mostly investigated the effects of SNPs in single SNP level and only a few studies considering the co-presence of multiple SNPs are available which showed differential effects on clinical manifestations of the disease^19–21^. Besides, only negligible focus was given on the role of HbE level in disease amelioration. In addition, till date there is no comprehensive study on the phenotypic variability and genetic modifiers of the diseases severity of HbE/β-thalassemia patients in Bangladesh.

With a view to gain more detailed understanding of the mechanism of clinical heterogeneity of HbE/β thalassemia, the present study aimed to highlight on several hematological and genetic factors including (1) comparison of HbF and HbE levels among mild, moderate, and severe groups, (2) identification of SNPs which could possess significant HbF-inducing effects on disease manifestations of Bangladeshi HbE/β thalassemia patients, (3) combined effect of the SNPs on HbF induction, (4) cumulative effect of multiple HbF inducing SNPs among mild, moderate and severe patients with c.79 G>A + IVS1_5G>C mutation covering 73.8% of the total patients (5) searching exceptional cases of mild, moderate, and severe cases and explanation of the mechanism.

## Methods

### Participant recruitment

The cross-sectional study performed on a total of 130 HbE/β-thalassemia patients (80 males, 50 females) and 50 unrelated healthy individuals over a period of 15 months from November 2018 to January 2020. Using a purposive sampling method, these patients (age range of 6 to 65 years) were enrolled from Bangladesh Thalassemia Samity Hospital located in Dhaka, where they came for follow-up examination and blood transfusion. The sample size was enumerated statistically appropriate for correlation study^22^.

A written informed consent form along with a structured questionnaire of information about the age of onset, age of first transfusion, transfusion interval, and splenectomy status of the patients were obtained prior to the enrollment in the study. The study was approved by National Ethics Review Committee (NERC) of Bangladesh Medical Research Council (BMRC).

### Hematological analysis and Hemoglobin study

Approximately 5.0 mL of venous blood was collected from the each participant just before blood transfusion and immediately transported to laboratory. Complete blood count (CBC) was performed on an automated hematology-analyzer (Sysmex Corporation, Kobe, Japan) to analyze hematological parameters. Hemoglobin variants were evaluated by Hb electrophoresis (Sebia CAPILLARYS2 Flex Piercing, Lisses, France).

### Mutation detection in β-globin and α-globin gene

Genomic DNA was extracted from whole blood using QIAGEN flexigene® DNA kit (Qiagen, Hilden, Germany) following manufacturer’s instructions. β-globin gene mutations using extracted DNA were detected using Real-time PCR-HRM analysis^23^. In α-globin gene, -α^3.7^ deletion and ααα^anti3.7^ triplication were detected by conventional Gap-PCR as per Liu Y et al^24^.

### Genotyping of eight SNPs in four major modifier genes

PCR-RFLP was used to identify rs7482144 (−158 Xmn1-Gγ polymorphism) using newly designed primer set and amplification conditions with necessary modifications of the procedure as described by Said et al^25^. 2μL of the purified PCR product was digested with 4 units of Xmn1 restriction enzyme (NEB, UK) following manufacturer’s instructions and electrophoresed on a 1.5% agarose gel.

Then, we established in-house Real-time PCR-HRM curve based methods for rest of the 7 SNPs-rs4895441, rs28384513, rs28384512 (HBS1L_MYB), rs11886868, rs4671393, rs766432 (BCL11A) and rs2071348 (HBBP1). A total of optimized 14 sets of primers (obtained from IDT, USA) for 7 SNPs of human BCL11A, HBBP1 and HBS1L_MYB intergenic region were selected for sequencing and Real-time PCR-HRM. Primer sequences and other information for the reactions of these 7 SNPs were given in **Table S1**. Firstly, reference samples (randomly selected) were prepared for each of the SNPs by Sanger sequencing of the polymorphic region on ABI PRISM310 Automated Sequencer (Applied Biosystems, USA). Then real-time PCR-HRM was performed on BioRad CFX96 Touch Real-Time System with SYBR® Green PCR Master Mix (Applied Biosystems). The thermal cycling profile was-initial denaturation at 95°C for 3 mins; 40 cycles of denaturation at 94°C for 10 seconds, primer annealing for 30 seconds at 63°C for HBBP1 and 60°C for BCL11A and HBS1L_MYB, extension at 72°C for 30 seconds, and then final extension at 72°C for 5 minutes. The subsequent melt curve was generated by 1min denaturation at 95°C, 1min renaturation at 60°C, and then melting at 60°C to 95°C with an increment of 0.1°C per 5 seconds. The target SNPs from unknown samples were identified by comparing with the clusters of reference samples.

### Statistical Analysis

Continuous variables were reported as mean±standard deviation (SD) and compared using the Mann–Whitney U-test and the Kruskal–Wallis one–way ANOVA test. Categorical variables were expressed as count and percentages. Hardy-Weinberg equilibrium was tested for each SNP distribution. Chi-square or Fisher’s exact tests were performed and odds ratios (ORs) with 95% confidence intervals (CIs) were calculated to compare the frequency of risk genotypes among different severity groups. Pearson’s correlation coefficient analysis was performed to study the association of HbF and HbE level with disease severity. The combined effect of SNPs on HbF level was analyzed by Tukey’s HSD post-hoc test. All these analyses were performed using GraphPad Prism®V.9.0 (GraphPad Softwares, Inc) where P<0.05 was considered statistically significant. To estimate the reduction of severity associated with the presence of favorable alleles, multinomial logistic regression analysis was performed by the Statistical Package for the Social Sciences software package (version 17.0; SPSS, Inc., Chicago, IL, USA). Odds ratios were estimated with a 95% CI.

## Result

### Demographic and baseline characteristics of the different severity groups of the patients

The study enrolled a total of 130 patients with the age range of 6 to 65 years (80 males and 50 females). As per Mahidol severity criteria, the patients were classified as (1) mild or Non-transfusion dependent (NTD) with ≤ 3 severity score (N=39), (2) moderately severe with 4-7 score (N=35), and (3) severe with **>**7 score (N=56). Patients of these three groups were similar in terms of age and gender distribution but differed in ages (months) of first blood transfusion and transfusion intervals (days), splenomegaly and splenectomy status. As expected, the patients in the severe group started 1^st^ transfusion at the earliest age (22.04 ±11.14 months) and required most frequent blood transfusion (TI=21.16±8.44 days) as compared to the moderate and mild group (AFT: 57.90±44.21 & 171.90±119.21 months, TI: 38.27±17.87 & 125.37±90.4 days respectively). Also, the rate of splenectomy and splenomegaly was much higher (17.8% and 41.07%) in the severe group than the moderately severe (11.4% and 34.28%) and mild (2.5% and 28.2%) patients (Table 1).

**Table 1:** Baseline information of the different groups of patients based on phenotypic severity classified according to Mahidol severity score* for HbE/β-thalassemia patients.

| Severity Groups | Mild / Non-transfusion dependent (score ≤ 3) | Moderate (score= 4-7) | Severe (score > 7) |
| --- | --- | --- | --- |
| Number of Patients, n (%); N= 130 | 39 (30%) | 35 (26.92%) | 56 (43.08%) |
| Male, n (%) | 23 (59%) | 21 (60%) | 36 (64%) |
| Female, n (%) | 16 (41%) | 14 (40%) | 20 (36%) |
| Age (Years)<br>Mean ± SD | 24.38 ± 11.83 | 17.30 ± 9.94 | 17.61 ± 7.49 |
| Age of 1<sup>st</sup> Transfusion (Months), Mean ± SD | 171.90 ± 119.21 | 57.90 ± 44.21 | 22.04 ± 11.14 |
| Transfusion Interval (Days), Mean ± SD | 125.37 ± 90.40 | 38.27 ± 17.87 | 21.16 ± 8.44 |
| Splenomegaly | 28.2% (11/39) | 34.28% (12/35) | 41.07% (23/56) |
| Splenectomized | 2.5% (1/39) | 11.4% (4/35) | 17.8% (10/56) |

\* Mahidol severity score includes 6 phenotypic clinical criteria, ranged between 0 (asymptomatic) to 10 (most severe)

### Role of fetal hemoglobin (HbF) and hemoglobin E (HbE) level in the disease manifestation HbE/β-thalassemia

Further we investigated the Hematological parameters and the Hemoglobin variants of the study participants to know their contribution on the variable disease severity. Among the hematological parameters, only HbE and HbF levels are almost exclusively of patient origin. The HbE levels (g/dL) of mild/NTD patients was signicantly higher (2.88±1.47) than moderate (1.36±1.1), and severe patients (1.35±0.97) which indicate a protective role of HbE against HbE/β thalassemia. Similarly, HbF levels (g/dL) of mild, moderate, and severe patients were 2.03±2.05, 0.84±1.27, and 0.58±0.78 **(Table S2)** and the differences in HbF among the 3 groups were significant (P<0.0001).

The study found a significant positive Pearson correlation of HbF and HbE concentrations with the age of first blood transfusion (p <0.0001/*r= 0.51* for HbF and p <0.005 and r= 0.28 for HbE) and transfusion intervals (p <0.0001/r= 0.56 for HbF and p <0.005/r=0.29 for HbE) **(Fig. A)**. On the other hand, both HbF and HbE concentrations displayed strong negative correlations with the clinical score of the patients (p<0.0001/r=-0.55 for HbF and p<005/r=-3.24 for HbE). Together, the findings indicate both HbF anf HbE are protective to Hb E/β thalassemia.

### Causative mutations in the β-globin gene among HbE/β-thalassemia patients in Bangladesh

A total of 11 different mutations were detected in the β-globin allele trans to HbE allele. IVS1_5G>C was the most common β-globin gene mutation and c.79G>A (HbE) + IVS1_5G>C combination could represent 73.8% of HbE/β-thalassemia patients distributed in all the three severity groups (Table 2). Among the 11 different mutation conbination; only two genotypes, namely c.79G>A+c.3G>T and c.79G>A + c.51delC + c.33C>A were exclusive for mild patients (n=3 for each genotype). However, the mutation patterns and corresponding distributions of patients among 3 different severity groups of HbE/β-thalassemia patients could not provide exclusive information to indicate whether certain mutation/combination of mutations in the β-globin allele were responsible for distinct clinical severity. Therefore, the study wanted to investigate the effects of other genetic modifiers to alter disease manifestations caused by HbE/β-thalassemia.

**Table 2.** Causative mutations in the β-globin gene allele trans to HbE allele among HbE/β-thalassemia patients in Bangladesh.

| β- genotype |  | % (n) | Phenotypic severity |  |  |
| --- | --- | --- | --- | --- | --- |
|  |  | N=130 | Mild/NTD (n); N= 39 | Moderate(n); N=35 | Severe (n); N=56 |
| 1 | c.79 G>A + IVS1_5G>C | 73.8 (96) | 23 | 24 | 49 |
| 2 | c.79 G>A + c.27_28 ins G | 1.5 (2) | 1 | 1 | 0 |
| 3 | c.79 G>A + c.126_129delCTTT | 5.4 (7) | 3 | 3 | 1 |
| 4 | c.79 G>A+ c.3G>T | 2.3 (3) | 3 | 0 | 0 |
| 5 | c.79 G>A + c.51delC + c.33C>A | 2.3 (3) | 3 | 0 | 0 |
| 6 | c.79 G>A + c.92 G>C | 3.1 (4) | 2 | 1 | 1 |
| 7 | c.79 G>A + 126delC | 1.5 (2) | 1 | 1 | 0 |
| 8 | c.79 G>A + IVS1_1 G>A | 3.8 (5) | 3 | 0 | 2 |
| 9 | c.79 G>A + IVS1_130 G>C | 2.3 (3) | 0 | 2 | 1 |
| 10 | c.79 G>A + c.46delT | 2.3 (3) | 0 | 2 | 1 |
| 11 | c.79 G>A + c.47G>A | 1.5 (2) | 0 | 1 | 1 |

### Disease-modifying effect of HbF inducing single nucleotide polymorphisms (SNPs)

Based on previous studies^10, 12, 26^, we targeted 8 SNPs in four major HbF Quantitative Trait Loci (QTL)-rs7482144 (−158 XmnI-Gγ) of HBG2 gene; rs4895441, rs28384513 and rs28384512 of HBS1L-MYB intergenic region, rs11886868, rs4671393, rs766432 on exon2 of BCL11A gene and rs2071348 on HBBP1 gene to detect the HbF inducing potential genetic modifiers of the disease severity (Table 3). The highest MAF (minor allele frequency) was found for HBS1L-MYB rs28384513 with the frequency of 0.46 followed by −158 XmnI-Gγ (MAF:0.45), HBS1L-MYB rs28384512 (MAF: 0.45) and rs2071348 in HBBP1 gene (MAF: 0.28) **(Table S3)**.

**Table 3.** Association of the 8 SNPs present in four genes with fetal hemoglobin level and the clinical severity score of HbE/β-thalassemia patients.

| Gene | SNP locus | Genotype (n) | Frequency among different severity groups %, (n) | | | Odd Ratio (95% CI);<br>$p$ value | HbF level (g/dL) | | clinical score | |
| --- | --- | --- | --- | --- | --- | --- | --- | --- | --- | --- |
| | | | Mild<br>n=39 | Moderate<br>n=35 | Severe<br>n=56 | | Mean | $p$ | Mean | $p$ |
| HBG2 | XmnI (C>T) | CC (21) | 2 | 5 | 14 | Reference | 0.49 | **0.0005** | 5.5 | **0.009** |
|  |  | CT (101) | 32 | 28 | 41 | **4.88** | 1.35 |  | 4.6 |  |
| | | TT (08) | 5 | 2 | 1 | **(2.1 -22.1)**<br>$p = 0.019$ | 1.84 | | 3.0 | |
| HBS1L-MYB | rs28384512 (C>T) | TT (38) | 14 | 8 | 16 | Reference | 0.98 | 0.07 | 4.6 | 0.95 |
|  |  | CT (67) | 15 | 18 | 34 | 0.68 | 1.5 |  | 4.3 |  |
| | | CC (25) | 10 | 9 | 6 | (0.31-1.53),<br>$P = 0.13$ | 1.2 | | 4.4 | |
|  | rs4895441 (A>G) | AA (95) | 21 | 24 | 50 | Reference | 1.01 | **0.0001** | 5.0 | **0.0005** |
|  |  | AG (25) | 13 | 08 | 04 | **2.72** | 1.58 |  | 3.6 |  |
| | | GG (10) | 05 | 03 | 02 | **(1.18-6.26)**<br>$P = 0.009$ | 2.58 | | 3.1 | |
|  | rs28384513 (T>G) | TT (33) | 08 | 07 | 18 | Reference | 0.91 | **0.0001** | 4.9 | 0.06 |
|  |  | TG (67) | 20 | 19 | 28 | 1.86 | 1.13 |  | 4.7 |  |
| | | GG (30) | 11 | 9 | 10 | (0.84-4.13)<br>$P = 0.06$ | 1.92 | | 3.9 | |
| BCL11A | rs11886868 (C>T) | CC (63) | 17 | 19 | 30 | Reference | 1.37 | 0.22 | 4.6 | 0.55 |
| | | CT+TT (67) | 22 | 16 | 26 | 1.51<br>(0.71-3.2)<br>$P = 0.14$ | 1.19 | | 4.4 | |
|  | rs4671393 (G>A) | GG (101) | 29 | 30 | 42 | Reference | 1.22 | 0.46 | 4.7 | 0.84 |
|  |  | AG (29) | 10 | 05 | 14 | 1.31<br>(0.54-3.15) | 1.4 |  | 4.5 |  |
| | | | | | | $P = 0.2$ | | | | |
|  | rs7664<br>32<br>(A>C) | AA (98) | 25 | 30 | 43 | Reference | 1.19 | 0.56 | 4.7 | 0.33 |
| | | AC (32) | 14 | 05 | 13 | 2.27<br>(0.98-5.2)<br>$P = 0.026$ | 1.45 | | 4.3 | |
| *HBBP1* | rs2071<br>348<br>(A>C) | AA (69) | 10 | 17 | 40 | Reference | 0.99 | **0.004** | 5.9 | **0.0001** |
|  |  | AC (44) | 20 | 13 | 12 | **4.35** | 1.26 |  | 3.9 |  |
|  |  | CC (17) | 09 | 05 | 04 | **(2.1- 9.2),**<br>**$P<0.0001$** | 2.42 |  | 3.7 |  |

Out of 8 SNPs, only 4 SNPs, namely −158 XmnI-Gγ, rs2071348, rs489544, and rs28384513 could show significant association with elevated levels of HbF in the study patients with HbE/β-thalassemia, generating p-values of 0.005, 0.0001, 0.0001, and 0.004, respectively. However, only the first 3 out of them were found associated with significantly reduced clinical severity scores (*p= 0.009, 0.0005, and 0.0001*, respectively). Also, Univariate analysis confirmed the links between these 3 predictors and severity showing a significant increased risk for causing mild clinical symptom in presence of the XmnI-Gγ SNP (OR 4.88 [2.1-22.1]), presence of G allele at HBS1L-MYB rs4895441 (OR: 2.72 [1.18-6.26]), presence of allele C at rs2071348 in HBBP1 (OR: 4.35 [2.1-9.2]). No significant association was observed with any of the 3 obseved SNPs in BCL11A or for the presence of rs28384513 and rs28384512 in the HBS1L-MYB inter-region. (**Table 3**).

### Combined effect of the 4 HbF inducing SNPs-rs7482144 (XmnI-Gγ), rs4895441(MYB-rs-48) & rs28384513 (MYB-rs-28), and rs2071348 (HBBP1) on raising HbF level and reduction of clinical severity of the HbE/β-thalassemia patients

To study the combined effect of the 4 HbF associated SNPs while they copresent in a same patient, we categorized the patients as XmnI(+) and XmnI(-) polymorphism and further sorted according to the presence/absence of other polymorphisms in HBS1L_MYB or HBBP1 gene. Then, Tukey’s test was used to compare the HbF levels among these groups of patients (**Figure 2**). HbF level was significantly increased in the patients with all different combinations of 3 different SNPs (MYB-rs-48, MYB-rs-28 and HBBP1-rs20) in the presence of XmnI-Gγ polymorphism.

**Figure 1.**
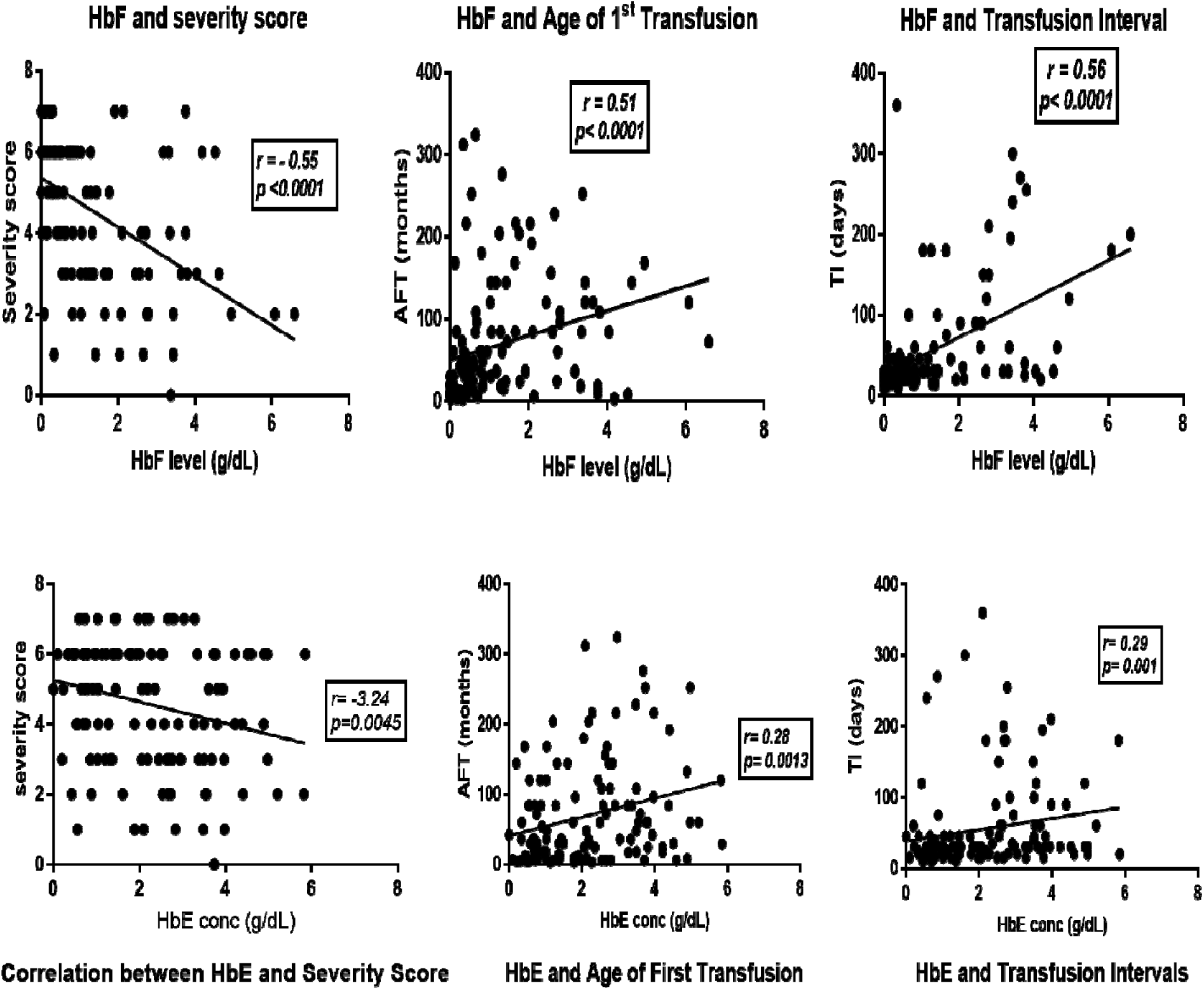
Pearson correlation analysis between HbF & HbE level and disease severity in patients with HbE/β-thalassemia. Disease severity was defined in terms of severity score, age of first transfusion in months (AFT)and transfusion intervals in days (TI).

**Figure 2.**
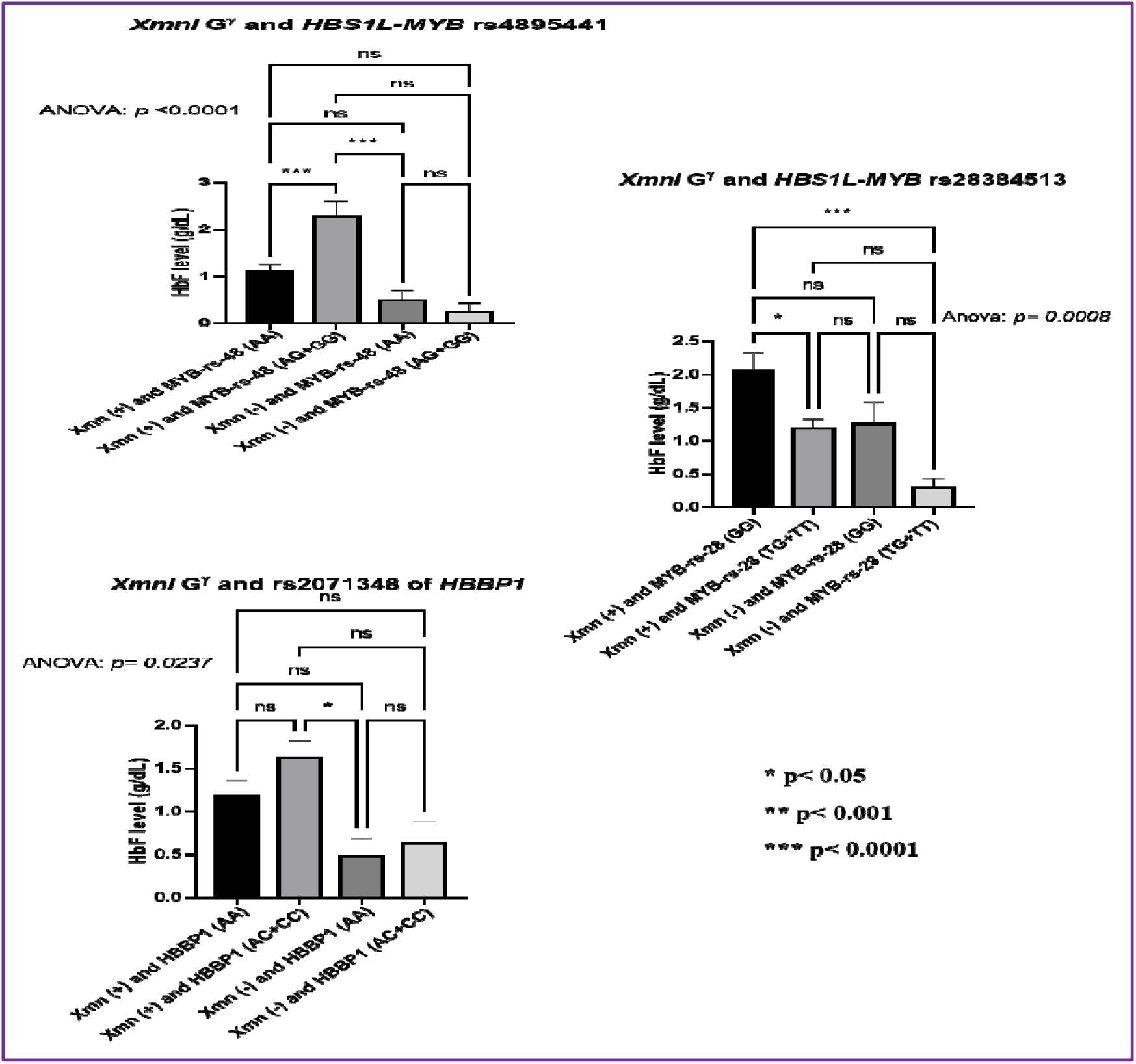
Comparison of HbF level for each combination of XmnI-^G^*γ* polymorphism with other 3 HbF associated SNPs-rs4895441, rs28384513 of HBS1L-MYB and rs2071348 of HBBP1 gene. The patients with co-presence of MYB-rs48 and the XmnI (+) polymorphism showed higher HbF level than the patients with corresponding XmnI (-) group. Similarly, for the rs28 in HBS1L_MYB, the XmnI (+) + GG combination showed the highest HbF level than the corresponding XmnI(-) + GG combination.. Similarly, the XmnI (+) + HBBP1(AC+CC) group had significantly highest HbF level.

Next, we aimed to detect the cumulative effects of these 4 SNPs on reducing severity in the HbE/β-thalassemia patients with c.79 G>A + IVS1_5G>C mutation combination covering 73.8% of the patients. Based on the results above, we defined T as the risk allele of XmnI; C allele for rs2071348, G for rs4895441 and rs28384513. To assess the cumulative effect of these loci, a multivariate analysis was performed using a logistic regression model. The SNP rs28384513 of HBS1L-MYB intergenic region was linked to severity with marginal significance (p=0.06) (Table 3). Mutinomoal Logistic Regression also showed that rs28384513 demonstrated insignificant association with reduced severity (OR=3.69, 95%CI: 0.62-7.52, *p=0.2*). Rest of the 3 favorable alleles were significantly associated with mild phenotype (Table 4). However, compared to individuals who had none of these risk variants, patients who carried any combination of 1, 2, 3 or 4 risk genotypes had a gradually increased risk (OR = 2.51, 5.47, 19.5, 39.0 respectively) of being less severe phenotype (Table 4).

**Table 4.** The analysis of cumulative effect of the 4 risk variants.

| Ameliorating SNP alleles |  | Number of patients |  | Odd Ratio | 95% CI<br>(*p value*) |
| --- | --- | --- | --- | --- | --- |
|  |  | Mild/NTD | Moderate + severe |  |  |
| Cumulative effect of the 4 risk variants | 0<br>(No SNP) | 1 | 13 | Reference |  |
|  | 1<br>(Single SNP present) | 6 | 31 | 2.51 | 0.3 - 23.02<br>(0.2) |
|  | 2<br>(any 2 SNPs present) | 8 | 19 | 5.47 | 0.7 - 49.16<br>(0.05) |
|  | 3<br>(any 3 SNPs present) | 6 | 4 | 19.5 | 2.6 - 145.5<br>(0.007) |
|  | 4<br>(all 4 SNPs present) | 6 | 2 | 39.0 | 4.4- 342.2<br>(0.001) |
| Multivariate logistic regression* | XmnI-<sup>G</sup><sub>γ</sub> (T allele) | 22 | 51 | 7.56 | 1.1- 71.05<br>(0.007) |
|  | rs4895441 (G allele)<br>in HBS1L-MYB | 13 | 5 | 4.67 | 1.25- 17.44<br>(0.02) |
|  | rs28384513 (G allele)<br>in HBS1L-MYB | 11 | 13 | 2.16 | 0.62 -7.52<br>(0.2) |
|  | rs2071348 (C allele)<br>in HBBP1 | 16 | 22 | 3.69 | 1.2 - 11.9<br>(0.03) |

NTD, Non-transfusion dependent; CI, Confidence Interval.
\* The reference category is 'severe'.

### Mechanism of clinical heterogeneity of HbE/β-thalassemia with c.79G>A+ IVS1_5G>C mutation

To explain the mechanism of diverse disease severity among the HbE/β-thalassemia patints with c.79G>A+IVS1_5G>C mutation including exceptional cases of mild, moderate, and severe cases, we also studied the effects of XmnI TT homozygosity and CT heterozygosity, coexistence of -α^3.7^ deletion, ααα^anti3.7^ triplication in α-globin chain along with the effect of variable number of SNPs present, which has been elaborated in Table 5. In some cases, HbE levels were taken into consideration to explain the disease manifestations.

**Table 5.**
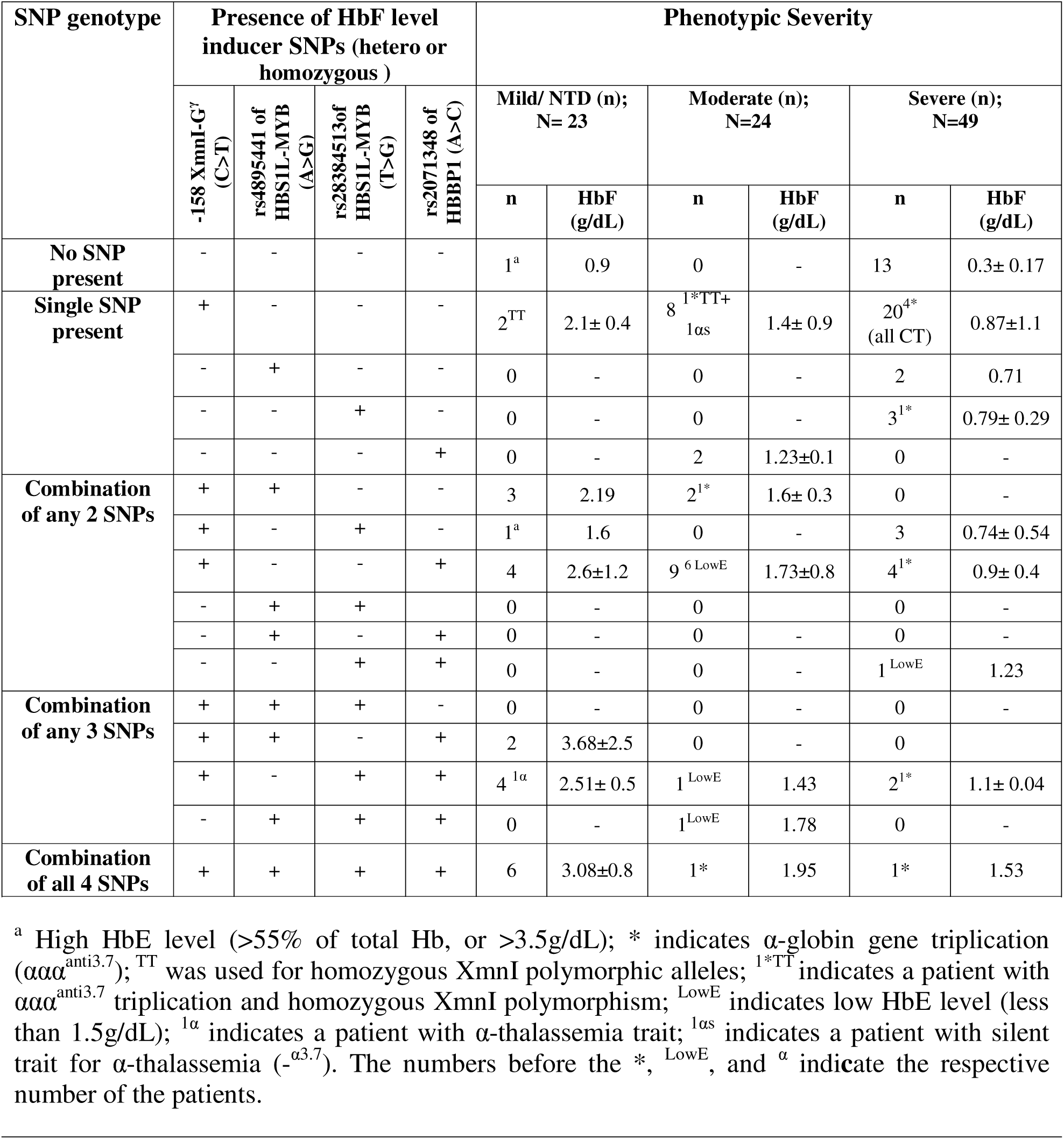
HbF inducing SNPs, HbF and HbE levels, and. α**-globin gene mutation(s) to explain the differential phenotypic severity of the 96 patients with c.79 G>A + IVS1_5 G>C mutation pair in the** β**-globin gene.**

In the absence of any SNPs, about 93% (13/14) of the HbE/β thalassemia patients (HbF=0.3± 0.17 g/dL) exhibited severe form of the disease. With presence of only −158 XmnI-Gγ, there were 66.7% (20/30, HbF= 0.87±1.1 g/dL) severe cases with CT heterozygosity and 26.7% (8/30, HbF=1.4± 0.9 g/dL) moderately severe cases (7 CT, only 1 TT which had α-globin gene triplication explaining the severity) and only 2 mild cases, both having XmnI TT homozygosity with increased HbF level of 2.1± 0.4 g/dL. Four of the 20 severe cases had also α-globin gene triplication (ααα^anti 3.7^) worsening the severity.

All the patients with rs489544 (n=2, HbF=0.71 g/dL) or rs28384513 (n=3, HbF= 0.79± 0.29 g/dL) in absence of XmnI polymorphism (rs7482144) were severe (one had α-globin gene triplication). Converesely, the patients with rs2071348 (n=2, HbF= 1.23±0.1) without Xmn1 had moderately severe disease.

For the co-presence of XmnI and rs4895441, 60% patients had mild/NTD thalassemia having markedly increased HbF (2.19 g/dL) and the rest 40% (n=2, HbF=1.6± 0.3 g/dL, one had α-globin gene triplication) had moderate form of the disease. While, co-presence of rs28384513 SNPs with XmnI, could not show the reducing effect on severity as 3 out of 4 patients were severe and notably, the only mild patient possessed very high HbE level (74%, 6.7 g/dL). Although rs2071348 of HBBP1 showed significant association with lowering the severity in both univariate and multivariate analysis, while coexist with XmnI, 55.6% patients (HbF=1.73±0.8 g/dL) were moderately severe as 6 out of 9 had low HbE levels. Similarly, some patients with combinations of 3 SNPs were severe or moderately severe due to having either low HbE level or coexistence of triplicated α-globin gene. Finally, out of 8 patients with co-presence of all 4 SNPs, six had mild form of the disease with markedly increased HbF (3.08±0.8 g/dL), while the rest 2 patients with α-globin gene triplication were severe and moderately severe despite of increased HbF (1.95 and1.53 g/dL respectively).

## Discussion

As a country on thalassemia belt, thalassemia has been found as a very concerning public health problem in Bangladesh with the expected annual birth of 7,193 HbE/β-thalassemia patients^27^. Unlike β-thalassemia major, the clinical manifestations of HbE/β-thalassemia are highly heterogeneous, mechanism of which is not clear enough^3^. The present study attempted to demonstrate the hematological and genetic basis of clinical heterogeneity of HbE/β-thalassemia by analyzing specimens of Bangladeshi patients. Without clearly knowing the mechanistic basis of the disease, management of such heterogeneous patients is quiet difficult for the physicians.

In this study, we first classifed the HbE/β-thalassemia patients according to the clinical severity^28^ and then studied the infuence of hemoglobin variants (HbF, HbE) along with the primary and secondary genetic modifers. Our study found that both HBF and HBE levels could contribute to protection against the disease and these findings are supported by the previous reports^29, 30^. The protective functions rendered by these two factors lies in the fact that HbF decreases the disease pathology by neutralizing excess unbound α globin chains in the nucleated erythrocytes and saving the cells from undergoing apoptosis and then subjecting them to differentiation to mature erythrocytes/reticulocytes^31^, whereas HbE functions by neutralization of unbound α-globin chain and transporting O_2_ to the tissues to fulfill O_2_ demand to some extent^16, 32^. Thus, a HbE/β-thalassemia patient with balanced production of HbE and HbF will have the mild form of the disease requiring no or occassional blood transfusion. Interestingly, we identified 5 mild patients having HbF levels (g/dL) of 0.5, 1.03, 0.8, 0.6, and 0.2 which were in the mean range for moderate and severe groups and were far below the mean level of mild patients. Remarkably, high levels of HbE (g/dL) (2.28, 3.24, 6.1, 3.78, and 2.26) were found for all these patients, indicating higher HbE levels were capable of making compensation for α-globin chain neutralization alongwith transporting O_2_ to the tissues^16^.

Types of mutations in HBB gene are the primary determinant of the disease severity in β-thalassemia^3^. A total of 11 different causative mutations in the β-globin allele trans to HbE allele were identified in the patients with HbE/β-thalassemia in Bangladesh. Two mutant genotypes, namely c.79G>A+c.3G>T and c.79G>A + c.51delC + c.33C>A were exclusively confined in mild patients (n=3 for each genotype), having 0% HbA for all. The first mutant group had both high HbE (avg 50.6%, 4.32g/dL) and HbF (avg 47.8%, 3.81g/dL) level thereby explaining the reason of the less severity (data not shown). The patients with c.79G>A+c.51delC+c.33C>A genotype had low HbF levels (avg 24.4%, 1.8g/dL) but significantly high HbE levels (avg 66.8%, 4.83g/dL) which explain why these patients were clinically mild and supported the finding in a recent study that the high-level HbE and the relatively low level of HbF together help the NTD patients to adapt to the anemia^1^. Similar to the neighbouring countries^33–35^ as well as other South Asian countries^36, 37^, IVS1_5G>C was the most common β-globin mutation and thus c.79G>A+IVS1_5G>C combination was found as most predominant mutation pair across all the 3 severity groups of patients covering 73.8% of HbE/β-thalassemia in the country. Therefore, the latter part of the study were mostly confined to patients with c.79G>A+IVS1_5G>C mutation combination to elaborate the mechanism of clinical heterogeneity of the disease. In this regard, the role of secondary modifiers like SNPs in 3 major HbF associated QTLs-1) HBG2 and HBBP1 gene in HBB locus, 2) HBS1L_MYB intergenic region, 3) repressor gene BCL11A and co-existence of α-thalassemia were considered which were reported to be associated with a milder clinical phenotype in HbE/β-thalassemia^3, 7, 12^. The highest minor allele frequency of 0.46 was found for HBS1L-MYB rs28384513 followed by Xmn1-^G^γ in HBG2 (0.45), HBS1L-MYB rs28384512 (0.45) while the lowest MAF (0.1) was found for the rs11886868 in BCL11A gene. For Xmn1-^G^γ polymorphism, nearly similar frequency was found in West Bengal and Malaysian population^38, 39^. However, frequency pattern observed for BCL11A and HBS1L_MYB SNPs was different from the previous report^40^.

Among the 8 SNPs studied, only 4 SNPs-Xmn1-Gγ, rs4895441, rs2071348, rs28384513 showed significant association with elevated level of HbF and the first 3 of them showed significant association with the reduced clinical scores of the patients with HbE/β-thalassemia. In our study, the strongest association in terms of both increasing HbF level and decreasing severity score, was observed with SNPs in the *HBG2* (Xmn1-Gγ) and *HBBP1* (rs2071348) gene in the HBB locus followed by rs4895441 and rs28384513 SNPs in *HBS1L*_*MYB* which was in line with previous findings of the study in India, Thailand and Indonesia^6, 10, 12, 41^.

Unlike most of the previous studies which discussed the disease-ameliorating role of HbF inducing SNPs in the disease-modifying genes individually^25, 42–44^, the present study also went further seeking the cumulative effects of multiple SNPs in changing the patterns of HBF induction as well as disease severity. Co-presence of multiple SNPs in different combinations changed the distribution patterns of HbE/β-thalassemia patients among mild, moderate, and severe groups. However, for any combinations of SNPs; mean HBF induction was highest in the mild group of patients and lowest in the severe group, whereas the moderately severe group had the HBF level in-between these two groups. Most of the patients (92.85%, 13/14) without any SNPs were severe and they could produce only very little HbF (<1 g/dL). Nevertheless, a patient without any SNP had mild form of the disease which could be explained by the presence of significantly high level of HbE (>3.5g/dL). Presence of only XmnI polymorphism failed to produce enough HbF and thus the patients of this category were either severe (20/30.) or moderately severe (8/30). Another reason for the high level severity of these patients is that they all had XmnI heterozogosity (CT) and 4 of them also had α-globin gene triplication worsening their condition. Only 2 patients of this category were mild having XmnI-homozygousity (TT) with 2.1±0.43g/dL HbF. Previous reports demonstrated that homozygous XmnI polymorphism could significantly increase HbF level with concomitant clinical improvement of splenomegaly and bone marrow expansion^45^. This is in agreement with our findings which showed that homozygosity of XmnI polymorphism could reduce the number of HbE/β-thalassemia patients with splenomegaly and splenectomy (data not shown).Therefore, we assume that one of the main reasons behind the very high number of trasfusion-dependent severe (56/130) and moderately severe (35/130) patients with HbE/β-thalassemia in Bangladesh might be due to the limited frequency of XmnI TT homozygosity (8/130) among the population.

By regression analysis, all 4 types of favorable allele were found to be signifcantly associated with mild clinical phenotype. XmnI (OR: 7.46) and HBS1L_MYB rs4895441(OR: 4.57) polymorphism were the most infuential modifers of the disease severity in our study population which is consistent with previous observations for β-thalassemia intermedia patients ^46, 47^.

In our study, it was observed that, 26.1% (6/23) of mild/NTD patients (HbF: 3.08±0.8g/dL) had inherited more than 3 ameliorating HbF boosting alleles as compared to only 1 (also had α-globin gene triplication) out of 49 patients (2.04%) in severe group. The data also indicates that all the patients with co-presence of all the 4 SNPs will have mild form of the disease provided that there is no α-globin gene triplication whereas with co-presence of 3 SNPs including XmnI polymorphism, 85.7% patients would have mild clicnical severity if there were enough HbE levels and no α-gene triplication (6 out of 7). Moreover, we found that the more risk genotypes of XmnI, rs28384513, rs4895441 and rs2071348 co-exist in a person, the less clinical severity in this person had, suggesting important links between these SNP variants in HBS1L-MYB intergenic region, γ-globin and HBBP1 along with a cumulative effect on both HbF level and severity score. Prominent link between XmnI in *HBG2* promoter and rs4895441 of HBS1L-MYB intergenic region has already been reported in few gene-gene interaction studies^21, 46^.

The main shortfalls of the study include (1) only 3 out of 12 known HbF inducing SNPs of BCL11A gene were analysed and none of them exhibited significant association with either HbF level or the severity score which contradicts several previous reports ^40, 43, 48^. (2) some other factors including KLF1 and tertiatry modifiers were not included in the study, and (3) detailed mechanism was discussed for the patients with c.79G>A+IVS1_5G>C mutation only but not for 10 other mutation combinations due to very low number of patients. Further studies on large samples are necessary to elaborate and validate the mechanism of clinical heterogeneity of HbE/β thalassemia.

Remarkably, this is the first study on the disease modifiers of HbE/beta-thalassemia in Bangladesh considering the effect of both primary and secondary level genetic modifiers of the disease severity. Despite a couple of shortfalls of the study, we believe that we could explain the hematological and genetic basis of clinical heterogeneity for majority of the patients with HbE/β thalassemia. The influence of SNPs can be different among different regions, reflecting a link between ethnicity and other genetic susceptibility factors. The findings of our study may assist the clinicians, to predict the clinical phenotype of HbE/β thalassemia at an early stage which may help in effective management of individual patient as well as revealing new therapeutic targets for increasing HbF levels in HbE/β-thalassemia patients in our population.

## Funding

This study was supported (in part) by research funding from Bangladesh Medical Research Council (BMRC), Dhaka, Bangladesh and under the Ph.D. Fellowship Funding from the Science and Technology Fellowship Trust, Ministry of Science and Technology, Government of Bangladesh.

## Acknowledgments

We sincerely thank all the patients and their families for their participation in this study, all the nurses, clinical staffs and the clinicians of Bangladesh Thalassemia Samity Hospital, Green Road Dhaka.

## Authorship

### Contribution

K.M conceptualized, planned and supervised the project, F.A.N and S.K.S designed the study; F.A.N. and M.H. performed the experiments; K.A., S.S.E., J.A.M., M.M.U.N., and G.S.V.assisted with the experiments; F.A.N., S.K.S and M.H. analyzed the data; N.S. and E.H. assisted in sample collection; F.Q. provided the funding and logistic support; F.A.N. and K.M. wrote the original manuscript; H.U.S. and W.A.K reviewed the manuscript.

### Conflict-of-interest disclosure

The authors declare no competing financial interests.

---

**Table S2:** Comparison of Hematological Parameters and Hemoglobin Variants among different severity groups.

| Parameters | NTD / Mild<br>N=39 | Moderate<br>N=35 | Severe<br>N=56 | ANOVA Test<br>(*p value*) | Healthy controls<br>N=50 |
| --- | --- | --- | --- | --- | --- |
| Total Hb (g/dl) | 7.78±1.23 | 7.08±1.44 | 6.92±1.03 | 0.0238 | 13.87±2.27 |
| RDW (%) | 28.05±8.5 | 24.24±8.87 | 22.97±7.59 | 0.0124 | 13.95±3.18 |
| MCV(fL) | 64.11±7.24 | 69.46±6.85 | 73.26±3.82 | 0.0001 | 81.72±4.57 |
| MCH(pg) | 20.21±2.9 | 21.99±3.08 | 23.28±2.27 | 0.0013 | 26.59±2.20 |
| Hb A (g/dL) | 2.27±2.81 | 3.89±2.42 | 5.33±2.61 | <0.0001 | 12.97±3.15 |
| HbE (g/dL) | 2.88±1.47 | 1.36±1.1 | 1.35±0.97 | <0.0001 | 0 |
| Hb F (g/dL) | 2.03±2.05 | 0.84±1.27 | 0.58±0.78 | <0.0001 | 0.15 ± 0.08 |
| Hb A2 (g/dL) | 0.33±0.1 | 0.27±0.1 | 0.23±0.07 | 0.0106 | 0.45 ± 0.46 |

Hb, Hemoglobin; MCV, Mean corpuscular volume; MCH, mean corpuscular hemoglobin; RDW, red cell distribution width; NTD, Non-transfusion dependent.
\*Results have been shown as mean ± SD, $p < 0.05$ was considered as significant.

## Notes

### Competing Interest Statement

The authors have declared no competing interest.

